# Remodeling of cholesterol metabolism in melanoma progression regulates nuclear membrane mechanics

**DOI:** 10.64898/2026.09.25.754527

**Authors:** Michelle A. Baird, Daniela A. Malide, Christian A. Combs, Robert S. Fischer, Clare M. Waterman

## Abstract

Metastatic cancer cells migrate through physically restrictive environments where nuclear deformation ruptures the nuclear envelope (NE) resulting in genomic instability. Although structural proteins of the NE are established regulators of nuclear mechanics, how nuclear membrane (NM) lipid composition contributes to its biophysical response to mechanical stress remains poorly understood. Here, we show that melanoma progression is associated with broad transcriptional remodeling of cholesterol homeostasis that is established early in disease progression, with altered expression of cholesterol biosynthesis genes associated with reduced patient survival. Building on our previous identification of the inner NM sterol reductase lamin B receptor (LBR) as a regulator of NE fragility, we find that LBR-dependent cholesterol biosynthesis promotes cholesterol enrichment within the NM and alters its spatial organization during cellular confinement. Quantitative fluorescence and lifetime imaging of biosensors of cholesterol distribution, lipid order, and tension reveals that cholesterol remodeling alters NM lipid organization and results in high NM tension in melanoma cells that is selectively reduced by cholesterol depletion. Cellular confinement generates nuclear blebs where cholesterol becomes preferentially enriched at highly curved, rupture-prone regions of the NM. Together, our results establish cholesterol-dependent NM remodeling as a regulator of nuclear mechanics and link metabolic changes acquired during cancer progression to the biophysical response of the NM to mechanical stress.

## Main Text

Metastatic cancer cells migrate through dense and mechanically restrictive tissue microenvironments where the nucleus, as the largest and stiffest organelle, becomes the primary physical and mechanical barrier to dissemination^1,2^. Excessive nuclear deformation can rupture the nuclear envelope (NE), transiently exposing chromatin to the cytoplasm and promoting DNA damage, genomic instability, and inflammatory signaling before nuclear compartmentalization is restored^3–8^. Chromatin organization, lamins, NE tethering proteins, actomyosin-generated forces, and mechanosensitive signaling pathways are established regulators of nuclear mechanics during deformation^9–19^. However, considerably less is known about how the nuclear membrane (NM) itself responds to mechanical confinement, or how its lipid composition contributes to its biophysical response to mechanical stress.

The NE is structurally composed of the inner and outer NMs (INM and ONM), nuclear pore complexes, membrane-associated tethering proteins, and the nuclear lamina. Together, the INM and ONM lipid bilayers form the NM, which maintains a specialized lipid composition that is required for nuclear organization and homeostasis^20–26^. The NM is enriched in unsaturated phospholipids, and contains relatively low levels of cholesterol and sphingomyelin, supporting the membrane fluidity required for nuclear pore complex insertion, membrane expansion during the cell cycle, and maintenance of nuclear architecture^27–33^. Additionally, reconstituted NM lipids in the absence of NE proteins and the lamina retain high fluidity and deformability, demonstrating that these mechanical properties are intrinsic to the lipid bilayers themselves^30^.

Consistent with this, perturbing NM lipid composition disrupts NE elasticity, curvature, organization, and integrity, suggesting that lipid composition directly contributes to NM biophysical properties^21–23,25,34,35^. Cholesterol represents a potentially important regulator of these properties, but its abundance, organization, and function within the NM remains poorly defined ^29,36^. In other cellular membranes, cholesterol is a major regulator of membrane thickness, order, fluidity, tension, bending rigidity, elasticity, and curvature ^37–42^. Whether cholesterol organization and abundance play similar roles in regulating NM mechanics remains unknown.

Our previous work identified lamin B receptor (LBR), an INM protein upregulated during melanoma progression, as a regulator of nuclear deformability and NE rupture during confined invasion^25^. LBR has dual functions at the NE, acting as a structural protein that tethers chromatin and lamins to the NE periphery, and also as a sterol reductase within the cholesterol biosynthetic pathway^43,44^. In confined melanoma cells high in LBR, we found NE rupture initiated with a local failure of the INM, followed by expansion of an ONM bleb and subsequent rupture, suggesting that local properties of the NM contribute to where mechanical failure occurs.

Importantly, LBR sterol reductase activity was required for increased nuclear deformability and NE rupture during confinement, directly linking cholesterol biosynthesis to nuclear mechanical responses in confinement^25^. LBR upregulation occurs early during melanoma progression, and remains elevated in advanced disease. This is particularly relevant in melanoma, which undergoes extensive metabolic remodeling during disease progression, including alterations in pathways regulating cholesterol^45–47^. Here, we investigated how cholesterol remodeling during melanoma progression alters the biophysical properties of the NM. We demonstrate that altered

cholesterol homeostasis changes NM cholesterol abundance and spatial organization, contributing to a higher tension NM state, and preferential cholesterol enrichment at highly curved, rupture-prone regions during confinement. Together, our findings link metabolic remodeling during melanoma progression to the biophysical response of the NM to mechanical stress.

## Results

### Melanoma progression is associated with broad remodeling of cholesterol homeostasis pathways

We hypothesized that the increase in cellular cholesterol observed during melanoma progression requires coordinated remodeling of cholesterol homeostasis pathways beyond the single biosynthetic step catalyzed by LBR. We therefore asked which cholesterol regulatory pathways are altered and when these changes occur during melanoma progression. We utilized annotated gene sets from the Molecular Signatures Database (MSigDB)^48,49^ to isolate genes involved in key aspects of cellular cholesterol homeostasis, including de novo cholesterol synthesis, transcriptional regulation, cholesterol uptake and trafficking, fatty acid synthesis, and indirect sterol regulation (Fig. 1A; Fig. S1A). Integration of these gene sets generated a panel of 72 cholesterol-associated genes that we used to examine transcriptional changes across melanoma progression (Fig. S1B,C). We utilized a previously established bioinformatics approach to analyze RNA-seq transcriptomic data from clinically staged, treatment naïve human melanoma lesions (GEO:GSE98394) that we previously subjected to differential expression analysis (DEA)^25,50,51^. This cohort is comprised of 27 benign nevi and 51 primary cutaneous melanomas representing the American Joint Committee on Cancer (AJCC) T1-T4 tumor thickness spectrum^52^. For this analysis, melanomas were grouped into early stage thin lesions (35; AJCC T1-T2) and late stage primary tumors (16; AJCC T3-T4), allowing us to determine when cholesterol associated transcriptional changes appear during primary melanoma progression.

**Figure 1.**
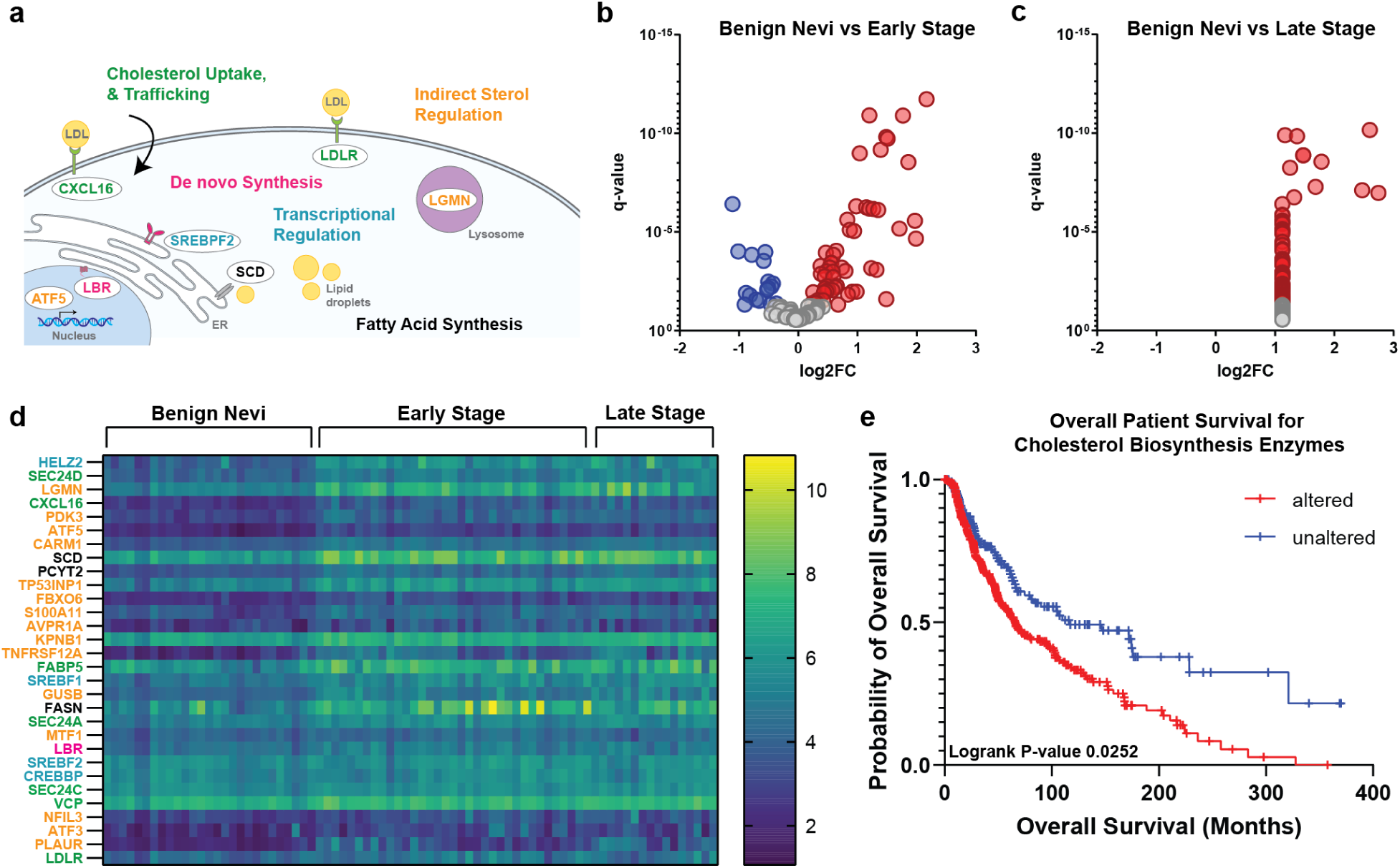
Melanoma progression is associated with broad remodeling of cholesterol homeostasis. **(A)** Schematic depicting functional pathways represented in a curated cholesterol homeostasis gene set (72 genes) including De novo Cholesterol Synthesis (Magenta); Transcriptional Regulation (Cyan); Cholesterol Uptake and Trafficking (Green); and Fatty Acid Synthesis (Black), and Indirect Sterol Regulation (Orange). Representative genes and associated cellular compartments are shown. **(B,C)** Volcano plots of differential expression analysis (DEA) of the 72 cholesterol-associated genes in RNA-seq data from clinically staged human lesions (GEO:GSE98394), comparing benign nevi (n=27) with early stage primary melanomas (n=35) (B) or benign nevi and late stage primary melanomas (n=16) (C). Differential expression is plotted as log2 fold change (log2FC) versus q-value. Red: Statistically significant up-regulation (red), and down-regulation (blue) are shown, non-significant genes with a q>0.05 (grey). **(D)** Heat map of the 30 cholesterol-associated genes exhibiting the largest fold changes between benign nevi and early stage primary melanomas across individual benign nevi, early, and late stage primary melanoma samples. Color scale indicates log2 fold change (log2FC). Genes were selected for visualization based on the fold change in the benign nevi versus early stage comparison and are ordered by fold change and color coded by functional pathway. **(E)** Kaplan-Meier analysis of overall survival from The Cancer Genome Atlas (TCGA) PanCancer Atlas melanoma cohort (SKCM) stratified by altered or unaltered expression of genes involved in de novo cholesterol biosynthesis (9 genes) identified as differentially expressed between benign nevi and early stage primary melanomas. (N=426 patients; Altered group=147, Unaltered group=279) Logrank P-value=0.0252. In (B.C) q ≤ 0.05 was used to determine significance. In (E) significance was determined by log rank test.

Comparison of benign nevi with early stage primary melanomas revealed broad upregulation of genes distributed across multiple functional pathways that contribute to cholesterol homeostasis, including enzymes involved in de novo biosynthesis (magenta; SQLE, LBR, TM7SF2), transcriptional regulation (cyan; SREBF2, SCAP, SREBF1), cholesterol uptake and trafficking (green; LDLR, LPL, CXCL16), fatty acid synthesis (black; SCD, FASN, ELOVL6), and indirect sterol regulation (orange; LGMN, PDK3, ATF5) (Fig. 1B; Fig. S1B). In contrast, comparison of early stage and late stage primary melanomas identified no significant differences among the cholesterol-associated genes examined (Fig. S1D), indicating that the major transcriptional changes in cholesterol homeostasis were already present in early stage melanoma and did not undergo further remodeling during progression to late stage primary disease. Comparison of benign nevi with late stage tumors further identified a subset of significantly altered cholesterol-associated genes, all of which were upregulated (Fig. 1C). To resolve the genes most strongly contributing to the early transcriptional response, we examined the 30 cholesterol-associated genes with the largest fold changes between benign nevi and early stage primary melanomas, identifying LBR as the only de novo cholesterol biosynthetic enzyme represented (Fig. 1D). Together, these data place early LBR upregulation within a broader program of cholesterol homeostasis remodeling that is established early and maintained throughout primary melanoma progression.

We next asked whether the cholesterol associated transcriptional changes identified during early melanoma progression was associated with patient outcome. For each functional cholesterol pathway, we assessed the subset of genes that were differentially expressed between benign nevi and early stage primary melanomas for their relationship with overall survival in The Cancer Genome Atlas (TCGA) PanCancer Atlas skin cutaneous melanoma (SKCM) cohort^53^. Altered expression of genes involved in transcriptional regulation, and uptake and trafficking were not significantly associated with overall survival (Fig. S1E-H). In contrast, altered expression of the de novo cholesterol biosynthesis genes upregulated during early melanoma progression was associated with a significant reduction in overall survival (Fig. 1E). Taken together, these data identify coordinated remodeling of cholesterol homeostasis as an early feature of melanoma progression and specifically link dysregulation of the de novo cholesterol biosynthesis program with poor clinical outcome.

### LBR-dependent cholesterol biosynthesis regulates cholesterol abundance and spatial organization within the NM

We next asked whether the transcriptional remodeling of cholesterol homeostasis during melanoma progression was accompanied by changes in cellular cholesterol distribution and whether this was altered during cellular confinement. To visualize steady-state cholesterol distribution at subcellular resolution, we performed super-resolution imaging of 1205Lu metastatic melanoma cells labeled with BODIPY-cholesterol (BD-Chol), a fluorescent cholesterol analog that enables visualization of relative cholesterol distribution within membranes^54,55^. Cells were co-labeled with SiR-DNA and mCherry-Sec61 to visualize chromatin and the ER/ONM, respectively^56,57^ (Fig. 2A). BD-Chol partitioned throughout the cell including at the plasma membrane (PM), lipid droplets, intracellular organelles, ER, and the NM, with line-scan analysis confirming overlay of BD-Chol and mCherry-Sec61 along the nuclear periphery (Fig. 2A, B; Fig. S2A). To account for variation in BD-Chol uptake and labeling, fluorescence within each membrane compartment was normalized to PM fluorescence within the same cell, which was selected as an internal reference based on its established high cholesterol content^58,59^. Under unconfined conditions, the ER and NM exhibited similar relative BD-Chol fluorescence, approximately 40% lower than the PM reference (Fig. 2C; Fig. S2B). We next asked whether LBR-dependent cholesterol biosynthesis regulates the relative distribution of cholesterol across intracellular membranes during confinement. Cells were confined to a height of 3 µm using a PDMS confinement device under lipid-depleted serum conditions to minimize exogenous cholesterol uptake, and BD-Chol distribution was compared between control and LBR-depleted cells (Fig. 2D). In comparison to 1205Lu cells, LBR depletion significantly reduced BD-Chol fluorescence in both the ER and NM relative to the PM (Fig. 2E), indicating that LBR-dependent cholesterol biosynthesis contributes to intracellular membrane cholesterol pools, including the NM.

**Figure 2.**
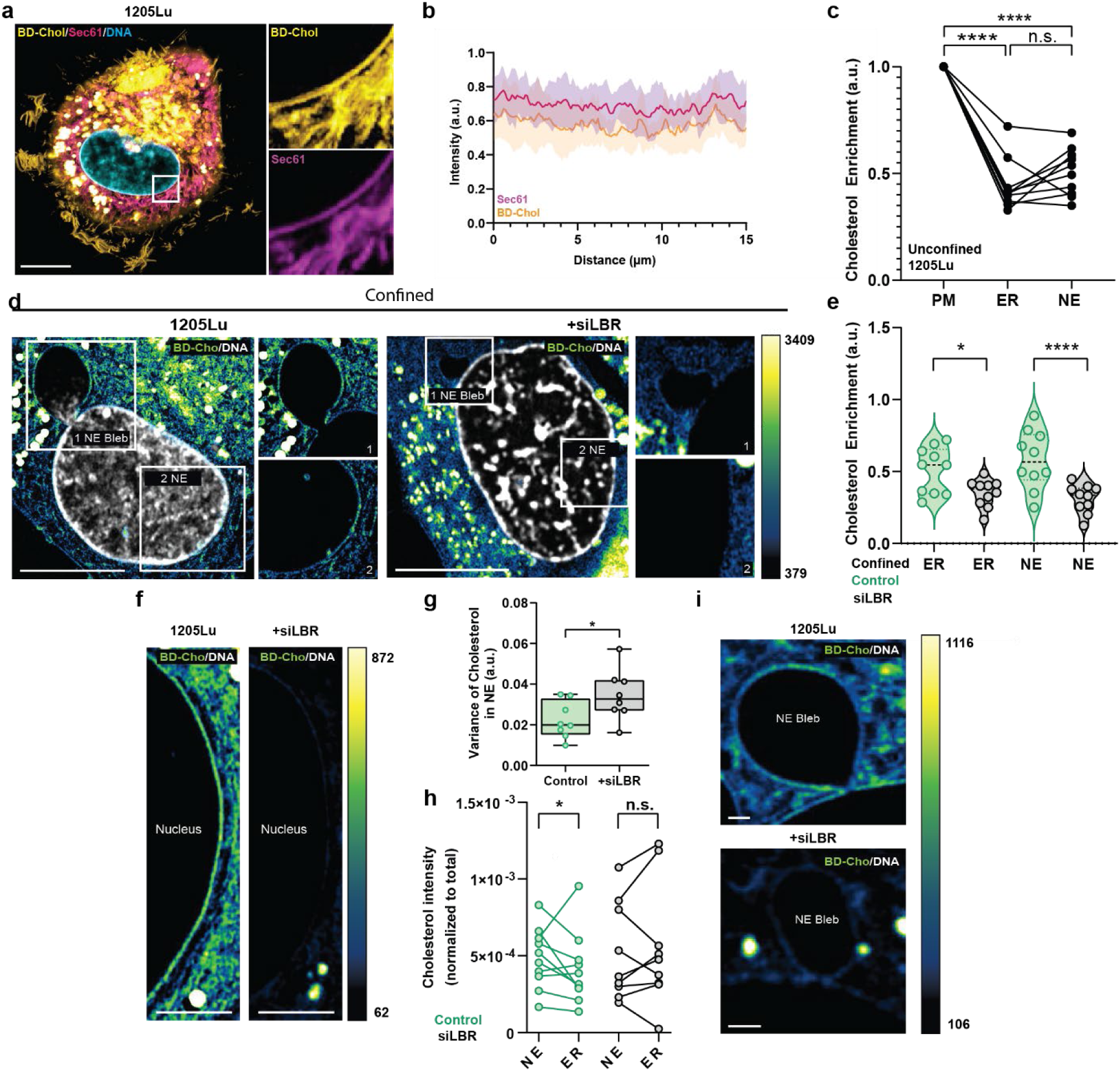
Cholesterol is spatially heterogenous within the nuclear membrane. **(A)** Representative super-resolution image of an unconfined 1205Lu melanoma cell labeled with BODIPY-Cholesterol (BD-Chol; yellow), mCherry-Sec61 (Magenta), and SiR-DNA (Cyan) to visualize cholesterol, the Endoplasmic reticulum (ER)/Outer nuclear membrane (ONM), and chromatin. Box indicate region enlarged at right. **(B)** Fluorescence intensity profile across 15 µm of the nuclear periphery showing spatial overlap of mCherry-Sec61 (Magenta) and BD-Chol (Yellow) at the NE (n=10 cells). **(C)** Quantification of normalized (to the plasma membrane (PM)) BD-Chol intensity measured for equal areas of the PM, NE and ER (n= 10 cells). From images similar to (A). Paired points represent measurements taken within the same cell. **(D)** Intensity color-coded super-resolution images taken during confinement to 3 µm of 1205Lu melanoma cells with (+siLBR, right) or without (1205Lu, left) transfection with siRNAs targeting LBR and stained with BD-Chol and SiR-DNA (grey). Numbered boxes indicate regions enlarged at right. **(E)** Quantification of normalized (to PM) BD-Chol intensity for NE and ER (D, n=10 cells) taken from images like those in (D) of confined cells. **(F, I)** Intensity color-encoded super-resolution images highlighting example NE and NE blebs taken during confinement to 3 µm of 1205Lu melanoma cells with (+siLBR, right) or without (1205Lu, left) transfection with siRNAs targeting LBR. **(G)** Quantification of variance in intensity measured along 2 µm long line scans of the NE taken from images of BD-Chol stained cells like those in (F). **(H)** Quantification of normalized (to total) BD-Chol intensity measured for equal areas of NE and ER (G, n= 3 experiments, 10 cells for control, 9 cells for LBR siRNA ). Paired points represent measurements taken within the same cell. (D-I) Experiments were performed in lipid depleted serum conditions, (A-C) were performed using complete media. Data points represent individual cells (C, E,G,H,I), or independent experiments each consisting of pooled cell data from each ROI (B). In (A, D) bar = 10 µm, in (F) bar= 5 µm, (I) bar= 1 µm. In (C,E) significance was tested with ANOVA, in (G,H), T-test with Welch’s correction.

We next asked whether LBR also regulates the spatial organization of cholesterol within the NM. Nuclear blebs generated during confinement represent highly deformed regions of the NM associated with subsequent NE rupture, suggesting that local differences in membrane composition could contribute to these mechanically vulnerable regions^25^. Control 1205Lu cells exhibited a relatively homogeneous distribution of BD-Chol along the nuclear periphery, with a slight enrichment in the NM compared to the ER (Fig. 2F-H). In contrast, LBR depletion produced regions of high and low BD-Chol fluorescence resulting in significantly greater variance in BD-Chol intensity along the NM (Fig. 2F-H; Fig. S2C). We therefore asked whether cholesterol was specifically enriched at nuclear blebs. Super-resolution microscopy identified BD-Chol enrichment at the base of nuclear blebs relative to adjacent non-bleb NM, and this regional enrichment was lost following LBR depletion (Fig. 2I; Fig. S2D). Together, these findings demonstrate that LBR-dependent cholesterol biosynthesis regulates both the abundance and spatial organization of cholesterol within the NM, promoting local cholesterol enrichment at highly deformed, rupture-prone regions during confinement.

### LBR-dependent cholesterol biosynthesis alters lipid organization in intracellular membranes in melanoma cells

We next asked whether the altered cholesterol distribution identified in melanoma cells was accompanied by changes in NM lipid organization. We utilized Laurdan (6-dodecanoyl-2-dimethylaminonaphthalene), a membrane-intercalating fluorescent vital dye with an emission spectrum highly sensitive to lipid order and composition, both of which alter membrane fluidity^60^. Laurdan excitation increases its movement in the membrane, causing dipole relaxation (DR) of the surrounding molecules and generating a shift in fluorescence lifetime and emission due to energy dissipation in the local membrane environment. Changes in lifetime of Laurdan’s blue emission are caused by water content in the membrane, where short lifetimes indicate water exclusion caused by increased lipid order, and longer lifetimes indicate water penetration allowed by lipid disorder^61^. Changes in lifetime of Laurdan’s green emission are caused by DR, where high cholesterol content in the bilayer results in a delay in the relaxation time. Exploiting fluorescence-lifetime imaging microscopy (FLIM) and phasor plot analysis, lipid order and DR can be separately evaluated independent of fluorescence intensity, on a pixel by pixel basis^62,63^. Spectral separation of these responses by FLIM and phasor plot analysis enables cholesterol-dependent membrane organization to be distinguished from changes in lipid order^62^. To independently analyze these properties, we combined FLIM imaging and phasor analysis of benign melanocytes, 1205Lu, and LBR KD cells stained with Laurdan (Fig. 3A-C; Fig. S3-S4).

**Figure 3.**
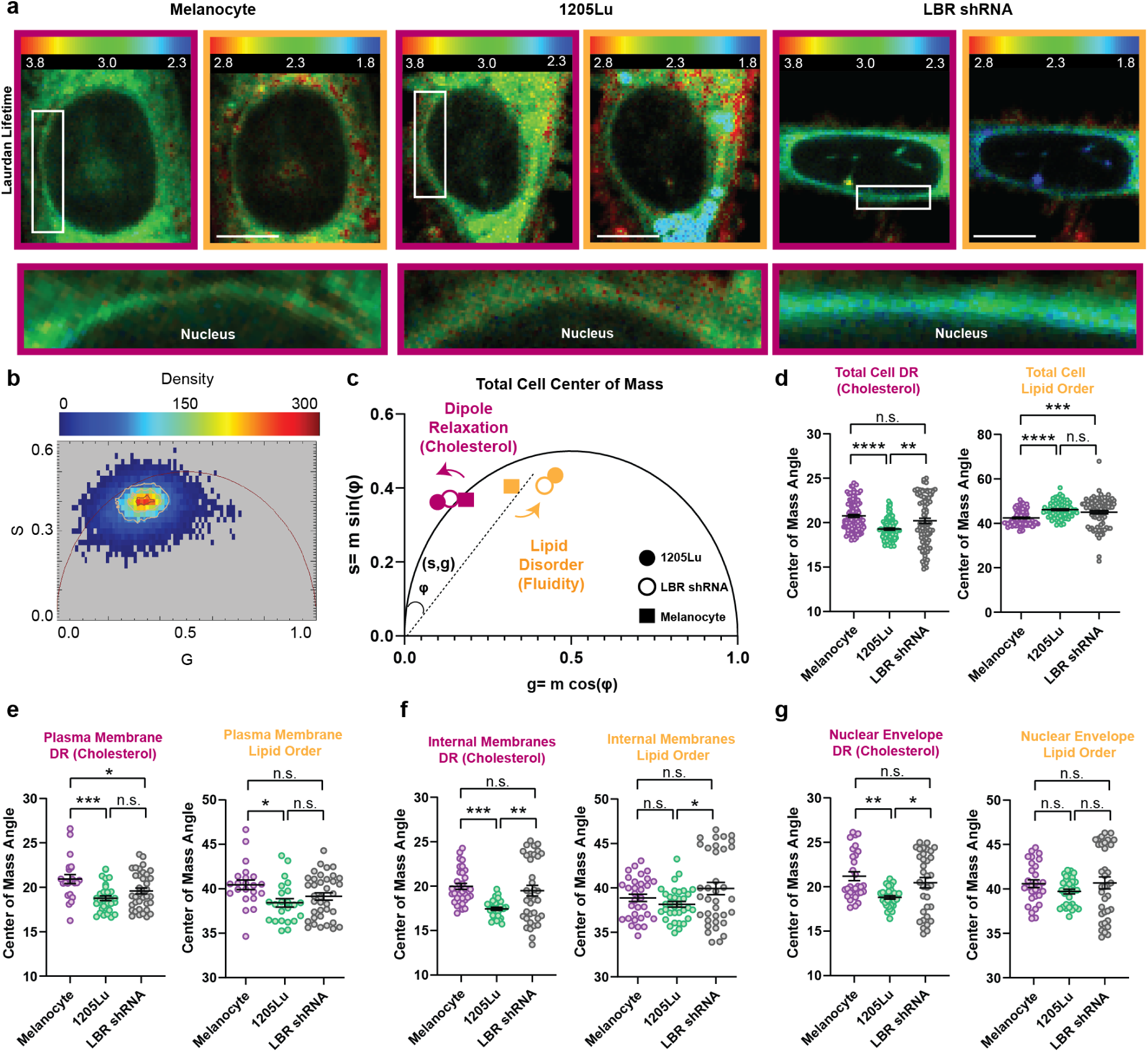
Melanoma progression alters plasma membrane lipid order and cholesterol-dependent dipolar relaxation in the nuclear membrane. **(A)** Representative FLIM images of unconfined Laurdan-stained melanocytes, 1205Lu, and 1205Lu cells depleted for LBR (LBR shRNA) in both the blue (orange boxes) and green (magenta boxes) emissions. Color scale represents FLIM lifetime values, box represents zoomed area along the NE (below). **(B,C)** Example phasor plot (melanocytes) and phasor analysis of Laurdan-stained melanocytes (square), 1205Lu (closed circle), and 1205Lu cells depleted for LBR (LBR shRNA; open circle). Orange data points represent the blue emission (lipid order), magenta data points represent green emission (dipole relaxation (DR)). Each point represents the center of mass of all pixel values on a phasor plot from a single field of view (FOV) containing multiple cells (similar to example in (b)). The angles between the centers of mass and the y-axis of the phasor plot are used to quantify the relative shift in Laurdan emission lifetimes for each experimental condition. **(D-G)** Quantification of angles of the centers of mass of all pixel values on a phasor plot from whole fields of views, each containing 3-15 cells in whole cells (WC) as well as segmented subcellular regions (plasma membrane (PM), internal membranes (IM), nuclear envelope (NE)) in melanocytes (WC: n=115 FOV, PM: n=22 FOV, IM: n=32 FOV, NE: n=29 FOV; N=3 independent experiments), 1205Lu (WC: n=81 FOV, PM: n=23 FOV, IM: n=34 FOV, NE: n=33 FOV; N=3 independent experiments), and 1205Lu cells depleted for LBR (LBR shRNA) (WC: n=87 FOV, PM: n=37 FOV, IM: n=36 FOV, NE: n=36 FOV; N=3 independent experiments). All experiments were performed in lipid depleted serum conditions. Data points represent individual FOV, each containing 3-15 cells. In (A) bar = 10 µm. Significance was tested with ANOVA and error bars = SD.

To substantiate our results using BD-Chol we performed phasor plot analysis of Laurdan’s lifetime in the green emission to determine if membranes in melanoma cells and melanocytes differed in their cholesterol-dependent membrane properties and the role of LBR in these differences. This analysis showed that 1205Lu cells exhibited a significantly delayed DR, consistent with increased cholesterol dependent membrane organization, compared to melanocytes, and this difference was lost upon LBR KD (Fig 3D-G; Fig. S3-S4). Analysis of subcellular regions showed that 1205Lu cells exhibited delayed DR and thus higher cholesterol in the PM, intracellular membranes, and NM compared to melanocytes. Consistent with the BD-Chol analysis, LBR depletion selectively caused a significant increase in DR within intracellular membranes and the NM, but not the PM, further supporting a preferential contribution of LBR to intracellular cholesterol pools (Fig 3D-G; Fig. S3-S4).

We then analyzed the fluorescence lifetime of Laurdan’s blue emission to determine whether melanoma progression was accompanied by changes in membrane hydration and lipid order. At the whole cell level, 1205Lu cells exhibited significantly longer Laurdan lifetimes than melanocytes, consistent with a more disordered, higher fluidity lipid environment (Fig 3D-G; Fig. S3-S4). To determine which membrane components contributed to this difference, we analyzed the PM, intracellular membranes, and the NE, identifying that the only significant difference in lipid disorder was at the PM (Fig 3D-G; Fig. S3-S4). Interestingly, cholesterol inhibition through LBR depletion had no effect on the lifetime in the blue emission, indicating that melanoma cells exhibit increased membrane fluidity predominantly through alterations in the PM, and this is independent of LBR-dependent cholesterol biosynthesis. Together these data indicate that melanoma progression is associated with an LBR-independent decrease in PM lipid order, while LBR promotes cholesterol and its effects on membrane organization in intracellular membranes, including the NM.

### Cholesterol-dependent remodeling increases NM tension and promotes nuclear rupture in melanoma

We next asked whether changes in cholesterol abundance and membrane organization identified in melanoma cells were accompanied by altered NM mechanical properties. To examine this, we utilized Flipper-TR, a mechanosensitive fluorescent membrane probe whose fluorescence lifetime reflects tension-dependent lipid packing^64^ (Fig. S5A). Lipid composition also influences this relationship, making Flipper-TR sensitive to the combined effects of membrane composition and tension^65^. Consistent with the high cholesterol content and lipid order of the PM, Flipper-TR lifetime was highest at the PM relative to intracellular membranes (Fig. S5B). Focusing on the NM, we compared NM Flipper-TR lifetime between benign melanocytes and metastatic 1205Lu cells. Under unconfined conditions, 1205Lu cells exhibited significantly longer Flipper-TR lifetimes at the NM than melanocytes (Fig. 4A-C), consistent with increased NM tension within the altered lipid environment of melanoma cells. To determine whether this difference was cholesterol dependent, cells were treated with MβCD. Cholesterol depletion significantly reduced NM Flipper-TR lifetime in 1205Lu cells toward levels observed in melanocytes, whereas melanocytes showed no significant response (Fig. 4A-C). Thus, elevated NM tension, as indicated by Flipper-TR lifetime, is cholesterol-dependent specifically in 1205Lus, consistent with cholesterol contributing to a higher tension NM state in melanoma.

**Figure 4.**
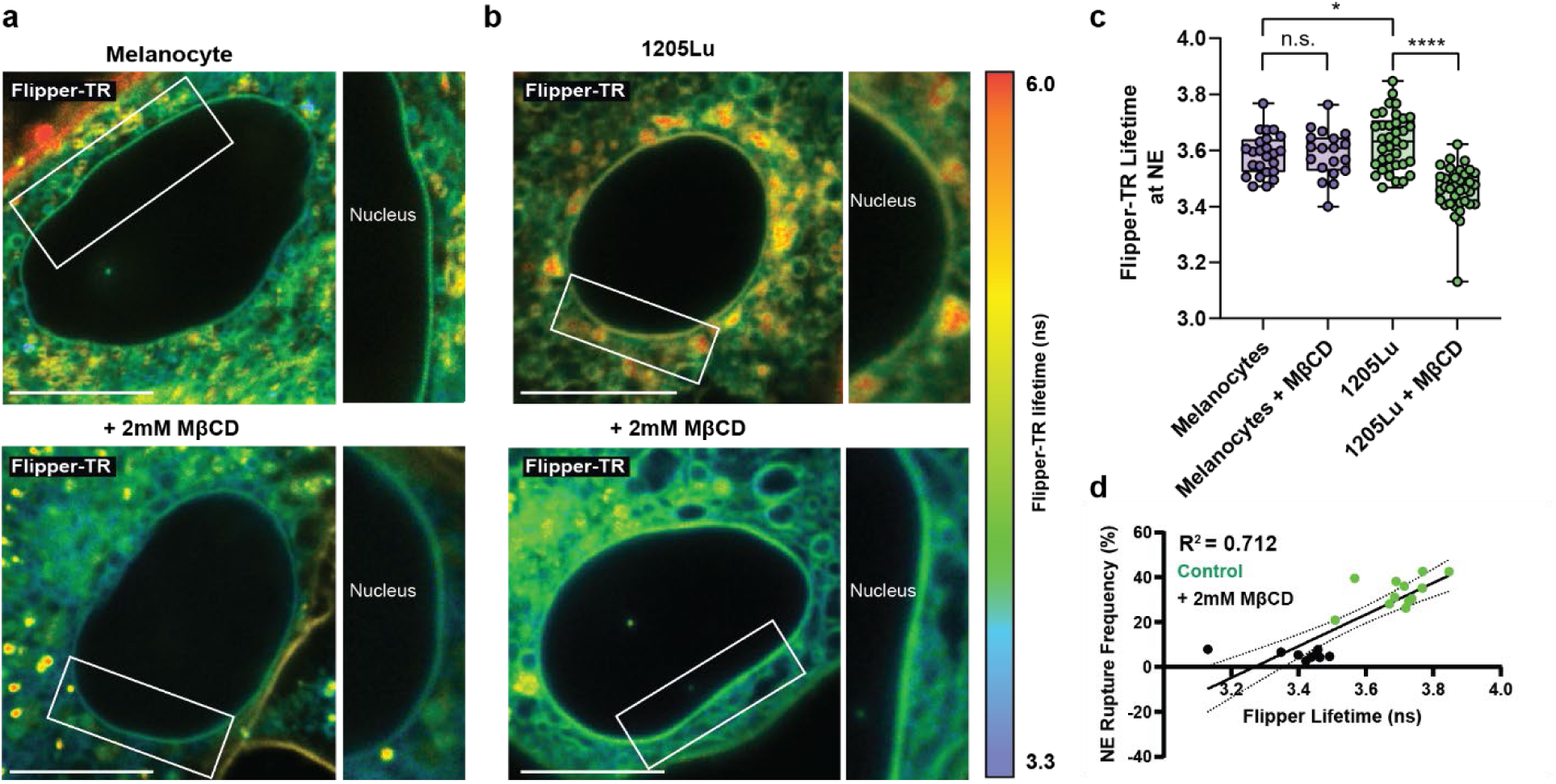
Cholesterol-dependent remodeling increases nuclear membrane tension. **(A,B)** Representative FLIM images of unconfined Flipper-TR labeled melanocytes and 1205Lu cells with (bottom) or without (top) treatment with 2 mM methyl-β-cyclodextrin (MβCD) (bottom). Color scale represents FLIM lifetime values, box represents zoomed area along the NE (right). **(C)** Quantification of the mean Flipper-TR fluorescence lifetime measured at the nuclear envelope (NE) in melanocytes and 1205Lu cells with or without cholesterol depletion with 2 mM MβCD. (1205Lu; n=37 cells, 1205Lu + MβCD; n=36 cells, Melanocytes; n=24 cells; Melanocytes + MβCD; n=19 cells. All conditions represent N=3 independent experiments. **(D)** Quantification of relationship between mean NM Flipper-TR lifetime and NE rupture frequency in 1205Lu cells with (+2 mM MβCD; black) or without (Control; green) cholesterol depletion with 2 mM MβCD (1205Lu; n=12 cells, 1205Lu + MβCD; n=9 cells. All conditions represent N=3 independent experiments.) Data points represent individual cells. In (A,B) bar= 10 µm. Significance tested with Student’s T-test (C). Error bars:mean; min and max. In (D) Significance was tested with simple linear regression, R^2^=0.7120.

Our previous work demonstrated that LBR-dependent cholesterol biosynthesis promotes NE rupture during mechanical confinement^25^, raising the question of whether this cholesterol-dependent NM mechanical state was associated with rupture susceptibility. We therefore examined the relationship between mean NM Flipper-TR lifetime and NE rupture frequency across experimental conditions. NE rupture frequency increased with mean NM Flipper-TR lifetime, with conditions exhibiting longer lifetimes also exhibiting greater rupture frequency (Fig. 4D). Together these findings suggest that cholesterol contributes to higher NM tension in melanoma, with the resulting mechanical state associated with increased NE rupture under confinement.

### Cholesterol organization is spatially coupled to NM curvature during confinement

We next asked whether the spatial organization of cholesterol within the NM was associated with specific local membrane geometries. Our earlier observation that BD-Chol was enriched at nuclear blebs during confinement suggested that cholesterol may preferentially accumulate in regions of extreme membrane curvature. To quantify NM curvature we used the ImageJ plugin Kappa to measure absolute curvature at every point along the nuclear perimeter^66^ (Fig. 5A). We first established the range of curvature states the NM could occupy under unconfined and confined conditions. In unconfined conditions, the NM was predominantly in a low-curvature state, with approximately half of the measured regions below 0.05 µm⁻¹, and the majority below 0.1 µm⁻¹ (Fig. 5B). Cellular confinement expanded this distribution to include higher curvature states, while LBR depletion shifted the population back to the low-curvature range which was characteristic of unconfined NM (Fig. 5B). Nuclear blebs were analyzed separately and occupied the highest curvature states, with curvatures of 0.5-1 µm⁻¹ distributed across the bleb membrane, and the most extreme curvatures of >2 µm⁻¹ concentrated at the bleb necks (Fig. 5A,B). Thus, confinement expands the NM beyond its basal low curvature range, into a broad distribution of curvature states, with the upper limits of this range localized to nuclear blebs and bleb necks.

**Figure 5.**
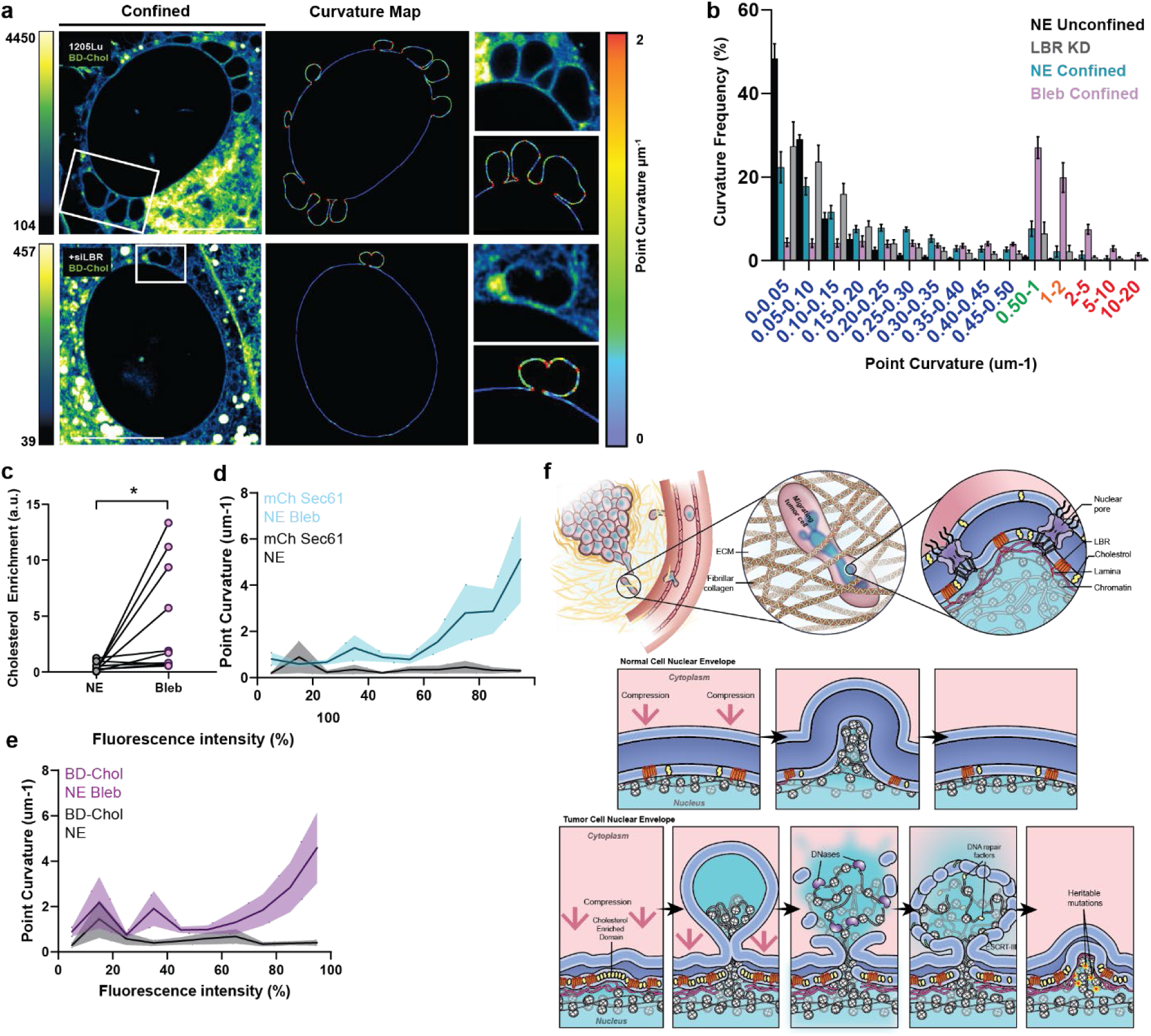
Cholesterol is preferentially enriched at high curvature regions of the nuclear membrane. **(A)** Representative super-resolution images of confined 1205Lu and LBR-depleted (+siLBR) 1205Lu cells labeled with BD-Chol and mCherry-Sec61, with corresponding nuclear membrane contours and point-curvature maps generated using Kappa. Boxes indicate regions enlarged at right. **(B)** Frequency distribution of absolute point curvature along the NM of unconfined 1205Lu cells, confined 1205Lu cells, confined nuclear blebs, and confined LBR-depleted cells. Curvature measurements were binned independently for each cell and plotted as the percentage of membrane positions within each curvature interval. (Cyan; 1205Lu Confined; n= 26,995 points, N=10 cells); (Black; 1205Lu Unconfined; n= 80,512 points, N=10 cells); (Purple; 1205Lu Confined NE Blebs; n= 55,809 points, N=10 cells); (Grey; 1205Lu LBR-depleted; n= 34,944 points, N=10 cells). X-axis colors reflect the point curvature LUT (A). **(C)** Paired comparison of BD-Chol enrichment within the highest 5% of fluorescence intensities (≥95th percentile) at nuclear blebs (Purple) and non-bleb NE regions (Grey) within the same cells. (N=10 cells). **(D,E)** Quantification of fluorescence intensity and local absolute point curvature within the NE and nuclear bleb regions. Fluorescence measurements were ranked independently within each cell and divided into 10 percentile bins, and mean absolute point curvature was calculated for membrane positions within each fluorescence percentile. Analysis was performed for mCherry-Sec61 (D) and BD-Chol (E) at the NE (Grey) or along NE blebs (Cyan); (N=10 cells). **(F)** Model depicting LBR-dependent cholesterol remodeling of the nuclear membrane during melanoma progression. Increased LBR-dependent cholesterol biosynthesis establishes a high cholesterol nuclear membrane that undergoes pronounced curvature and blebbing during cellular confinement, with preferential cholesterol enrichment at highly curved, rupture-prone membrane regions. Data points represent individual cells (C). In (A) bar= 10 µm. Significance tested with Student’s T-test (C). in (B) error bars = mean, SEM, (D,E) Colored error bars = SEM.

We next asked how BD-Chol fluorescent intensity was distributed across this expanded range of NM curvatures. Pairwise comparison of the highest 5% of BD-Chol fluorescence within individual nuclei demonstrated significantly greater cholesterol enrichment at nuclear blebs than at non-bleb regions of the NE (Fig. 5C). We next determined how this local cholesterol enrichment was related to membrane curvature across the bleb. Using an intensity based approach developed to relate local molecular enrichment to membrane curvature^67^, BD-Chol fluorescent intensity along the NM was ranked into percentile bins within individual cells, and compared with the corresponding local membrane curvature. To establish how membrane associated fluorescence varied with curvature independently of cholesterol, we first performed this analysis using mCherry-Sec61. Within nuclear blebs, mCherry-Sec61 fluorescence increased gradually with local curvature at higher fluorescence percentiles (Fig 5D). BD-Chol exhibited a distinct distribution, including intensity peaks at approximately 3.5 µm⁻¹, that were not observed with Sec61, correlating with the curvatures associated with nuclear blebs, followed by the highest BD-Chol intensities at the largest curvatures (Fig. 5E). In contrast, neither signal showed a strong curvature dependent distribution along non-bleb regions of the NE (Fig. 5D,E). Together, these findings demonstrate that cholesterol is preferentially enriched within highly curved nuclear blebs during confinement, connecting local cholesterol organization to regions of extreme NM geometry associated with NE rupture. Collectively, our data demonstrate that cholesterol dependent remodeling alters NM organization and mechanics in melanoma cells, establishing a high tension membrane state that is further spatially reorganized under confinement, to concentrate cholesterol at highly curved, rupture-prone regions (Fig. 5F).

## Discussion

Here, we show that transcriptional remodeling of cholesterol homeostasis occurs early during melanoma progression and is associated with changes in the organization and mechanical properties of the NM. Transcriptomic analysis of clinically staged primary melanomas revealed broad upregulation of pathways controlling cholesterol homeostasis early in disease progression, with altered expression of de novo cholesterol biosynthesis genes associated with reduced patient survival. Focusing on how this metabolic remodeling changes cellular cholesterol organization, super-resolution imaging of BD-Chol revealed that LBR-dependent cholesterol biosynthesis preferentially contributes to intracellular membrane cholesterol pools, including the NM, where it regulates both cholesterol abundance and spatial distribution. Using complementary fluorescence lifetime measurements of probes of lipid organization and membrane mechanics, we further demonstrate that melanoma cells exhibit elevated NM Flipper-TR lifetime that is cholesterol dependent and consistent with a high tension NM state. While under cellular confinement, cholesterol became preferentially enriched at highly curved nuclear blebs associated with NE rupture. Together, these findings connect altered cholesterol metabolism during melanoma progression with changes in NM organization and mechanics, identifying cholesterol as a link between metabolic state and the response of the NM to confinement.

How does cholesterol regulate the mechanical properties of the NM? Cholesterol is an established regulator of membrane organization and mechanics, yet its function within the NM remains poorly understood. Cholesterol is present at low levels within nuclear membranes, and the physical continuity of the ONM with the ER raises the question of how differences in lipid composition can be established and maintained across these connected membrane structures^68^. However, recent studies show that the lipid distribution between the ER and NE is not necessarily uniform, and have identified several lipid metabolism enzymes at the INM, supporting the possibility of local regulation of NM lipid composition^20,24,69–72^. Our findings identify LBR as one mechanism contributing to this regulation, where its sterol reductase activity controls the relative abundance and spatial organization of cholesterol within the NM. Importantly, cholesterol depletion reduced NM Flipper-TR lifetime specifically in melanoma cells, supporting a cholesterol dependent contribution to the mechanical state of the NM. We propose that lipid composition provides a dynamic layer of NM mechanical regulation alongside the structural contributions of lamins and chromatin. Redistribution of lipids within the bilayer could allow NM mechanics to be modified locally and rapidly without requiring large scale remodeling of structural networks and proteins, increasing nuclear adaptability during mechanical confinement.

How does the spatial organization of cholesterol alter the response of the NM to mechanical stress? We found that nuclear blebs generated during confinement contain the highest curvature regions of the NM, where cholesterol becomes preferentially enriched. The spatial relationship between cholesterol and curvature suggests that local lipid composition contributes to the organization of highly deformed membrane regions. Whether cholesterol promotes membrane bending, or curvature drives local cholesterol accumulation, remain open questions, and these processes may act together during nuclear deformation. Lipid composition directly influences the energetic cost of membrane bending, and reconstituted NM lipids containing cholesterol exhibit unusually high deformability, supporting the role of NM lipids and cholesterol in regulating membrane geometry^30,73^. Importantly, these cholesterol-enriched, highly curved regions corresponded to NM blebs where we previously demonstrated that NE rupture initiates, through local failure of the INM, followed by expansion and rupture of a bleb in the ONM. Increasing NM Flipper-TR lifetime was also associated with increasing rupture frequency across experimental conditions, connecting high NM tension with rupture susceptibility. We propose that cholesterol regulates how the NM tolerates mechanical stress during confinement. A membrane already maintained at a high tension state has less capacity to accommodate additional deformation without increasing local membrane stress. In melanoma cells, cholesterol contributes to a higher tension NM state, while confinement additionally concentrates cholesterol at highly curved, rupture-prone blebs. Thus, cholesterol may influence NM response to confinement at two scales, by contributing to its global mechanical state, and by becoming locally reorganized at regions of extreme membrane curvature. The merging of these properties at nuclear blebs provides a potential mechanism for spatially restricting NE rupture.

Our finding that cholesterol regulates NM mechanics identifies a biophysical consequence of the metabolic remodeling that occurs during melanoma progression. Cholesterol metabolism in cancer is primarily considered through its roles in membrane production, signaling, proliferation, and adaptation to metabolic stress^74^. Our findings identify an additional role for lipid metabolism in controlling how the NM responds to mechanical stress during metastatic invasion. This is particularly relevant given the experimental and epidemiological studies associating statin use with reduced melanoma progression or metastatic burden^75–78^, raising the possibility that altered membrane mechanics may represent an additional consequence of targeting cholesterol metabolism. Our patient data further demonstrate that cholesterol homeostasis is remodeled early during melanoma progression, suggesting that metabolic changes capable of altering NM properties are established before invasion and precede the extensive nuclear deformation required for migration through restrictive tissue environments. Together, our findings establish nuclear membrane lipid remodeling as a mechanism linking cancer metabolism to nuclear mechanobiology, demonstrating how metabolic changes acquired during tumor development can alter the physical response of cancer cells to the mechanical stresses of metastatic dissemination.

## Acknowledgements

We thank the Waterman Lab, Dr. Mehdi Pirooznia, Dr. Shirin Bahmanyar, Dr. Michelle Digman and Dr. Alexander Sodt for their feedback and discussions on this work, as well as the NHLBI light microscopy, bioinformatics, and DNA sequencing and genomics.

## Author contributions

Conceptualization: MAB, CMW, RSF Investigation: MAB, RSF, DM Supervision: CMW

Writing – original draft: MAB, CMW

## Data and materials availability

All data, plasmids, and sequencing data are available upon request.

## Supplementary Materials

Materials and Methods

Figs. S1 to S5

