## Supplementary Materials for "Remodeling of cholesterol metabolism in melanoma progression regulates nuclear membrane mechanics"

**The file includes:**

Materials and Methods
Figs. S1 to S5
References (1-12)

**Materials and Methods**

**Cell Culture**

1205Lu cells (Dr. Glenn Merlino, NCI) were cultured in Tu media (80% MCDB153, 20% L-15) supplemented with 2% fetal bovine serum (FBS; Atlanta Biologicals S11150), 2.5 ng/ml insulin (Sigma), and 1.68 mm $CaCl_2$. Immortalized human melanocytes (Dr. Glenn Merlino, NCI) were cultured in dermal cell basal media (ATCC, PCS-200-030), supplemented with the melanocyte growth kit (ATCC, PCS-200-041). All cells were maintained at 37°C at 5% $CO_2$ for <15 passages. For experiments examining the contribution of LBR-dependent cholesterol biosynthesis, 1205Lu cells were maintained for at least 7 days in media supplemented with 2% lipid-depleted FBS (Omega Scientific, Fisher Scientific) to minimize the contribution of exogenous cholesterol uptake. Transient expression of cDNAs or siRNAs was performed using the Amaxa nucleofector kit R (Lonza) and Amaxa Nucleofector II (Lonza) programs X-001 (1205Lu), U-024 (melanocytes) and 1.0 µg cDNA, 500nM for siRNA was used for $2x10^6$ cells.

**CDNA expression vectors and lentiviral expression**

mCherry Sec61 cDNA was used for transient transfections (Mike Davidson; Florida State University, Tallahassee, FL) (Addgene #55129). ON-TARGETplus SMARTpool siRNA (Dharmacon) was used for human LBR (J-021505-05), and cells were transiently transfected for 48 hours prior to experimental use. Generation of a custom LBR shRNA lentivirus plasmid was done using the 5'UTR duplex (5'GUAAAGGAGUGCUGUCUUAUU 3') RNA sequence to design a shRNA ligated into the pLKO-Tet-On lentivirus AgeI-EcoRI sites for expression under the control of the H1/TO promoter 1. 1205Lu cells stably expressing LBR shRNA (Dharmacon) were generated using the second generation lentiviral packaging plasmid psPax2 (Addgene 12260) and pMD2.G (Addgene 12259). Lentiviral particles were generated by first transfecting 1 million HEK293FT with Lipofectamine 2000 (Thermo Fisher) according to manufacturer's protocols using 2.5 µg plasmid DNA, 1 µg pMD2.G, and 2.5 µg psPax2 and collecting the virus-containing supernatant at 48 hours. Supernatant was then filtered through a 0.45 µm filter (Sigma), then immediately placed on target 1205Lu cells that had been seeded 24 hours before transduction at a 1:1 ratio of viral supernatant to 2% Tu media and allowed to incubate for 48 hours, upon which cells were passaged into media containing a final concentration of 2.0 µg/ml puromycin for selection. Knockdown was induced in cell culture using 100 nM doxycycline (Sigma) for 72 hours prior to experiments.

**Drug Treatment**

Acute depletion of cellular cholesterol was performed with Methyl-β-cyclodextrin (MβCD, Sigma) at a final concentration of 2 mM dissolved in water as previously described[1,2]. Cells were treated for 1 hour at 37°C prior to imaging, washed with PBS, and replaced with fresh media for data acquisition.

**BODIPY-Cholesterol Labeling**

To visualize the subcellular distribution of cholesterol, cells were labeled with BODIPY-cholesterol (BD-Chol; TopFluor-Cholesterol, Avanti Polar Lipids). To generate labeling solution, BD-Chol was dissolved at 1 mg/ml in a 1:1 (v:v) chloroform:methanol solution and dried under nitrogen gas. This was then complexed with 370 mM methyl-β-cyclodextrin (MβCD, Sigma) in PBS at a 100:1 (v:v) ratio, vortexed, sonicated in a water bath for 3 minutes, and incubated on a rocker overnight at 37°C. Undissolved dye was then removed by centrifugation at 16,000 x g for 15 min. 1205Lu cells were incubated with the BD-Chol/MβCD solution at 1:1000 dilution for 10 minutes, washed with PBS, then incubated in fresh media containing Blue-White DPX ER-tracker (to label the endoplasmic reticulum (ER)/outer nuclear membrane (ONM), 1 µM; Thermo) and SiR-DNA (to label the chromatin, 1 µM; Cytoskeleton) for 1 hour to allow for cholesterol internalization to cellular membranes and uptake of vital dyes.

**Laurdan Labeling**
To assess the lipid properties of cellular membranes, the dye laurdan (6-dodecanoyl-2-dimethylamino naphthalene; Sigma) was used. Prior to imaging, cells were cultured for 7 days in Tu 2% media (1205Lu) or dermal cell basal media (melanocytes) supplemented with lipid depleted FBS, LBR knockdown was induced with doxycycline 72 hours prior to imaging. On the day of imaging, a fresh stock of laurdan was prepared to prevent degradation of the probe. First, it was dissolved in 37°C DMSO to create a 5 mM stock solution, then further diluted to 50 µM in Opti-MEM (Gibco). Cells were incubated with Laurdan for 5 hours at 37°C to allow for cellular membrane dye uptake. Cells were imaged within 1 hour following the 5 hour labeling period to allow for consistent probe exposure between experimental repeats.

**Flipper-TR Labeling**
A 1 mM stock solution of Flipper-TR (Flipper-TR; Spirochrome) was prepared per manufacturer's instructions. This stock solution was diluted to 1 µM for imaging in serum-free Tu media that was prepared as described above. Cells were labeled for 1 hour at 37°C to allow for dye to label the internal membranes, washed with PBS, and replaced with fresh label free media for imaging.

**Bioinformatics**
***Data Acquisition and Differential Expression Analysis***
Patient bulk melanoma tumor RNA-seq data was downloaded from the Gene Expression Omnibus (GEO:GSE98394). The patient cohort was categorized based on the provided clinical staging information into benign nevi (27 samples), and 51 treatment naïve primary cutaneous melanomas grouped into early stage (35 tumors), and late stage (16 tumors) for differential expression analysis (DEA)[3]. Clinical staging information was based on the American Joint Committee on Cancer (AJCC). Primary melanomas were grouped as "early stage" (T1-T2) or "late stage" (T3-T4) for downstream analysis. RNA-seq data was subjected to quality control analysis using FastQC (v0.11.8)[4] . This was followed by alignment to the human genome (GRCh38/hg38) using spliced transcripts aligned to a reference (STAR)[5]. Gene expression was quantified using featureCounts (v2.0.0)[6]. RNAseq sample quality was determined using ESTIMATE (v 1.0.13) R package. Tumor purity estimates and sex were used as the covariates for DEA, performed using limma-voom [7]. All RNA-seq data was filtered to remove genes with an expression count < 5 in 90% of the samples were excluded before DEA. Differentially expressed genes were defined using FDR-adjusted P< 0.05 (q-value), to remove low-expressing and non-significant genes.
***Identification of genes of interest***
Cholesterol-associated genes were curated from Molecular Signatures Database (MSigDB)[8] Reactome_cholesterol_biosynthesis and Reactome_regulation of cholesterol by srebp databases and further classified into functional groups using published data characterizing their functional role. These genes include: De Novo Cholesterol Biosynthesis (FDFT1, ACSS2, LBR, IDI2, IDI1, SQLE, HSD17B7, TM7SF2, DHCR24); Transcriptional Regulation (SREBF2, SCAP, SREBF1, INSIG1, INSIG2, NFYB, CREBBP, MED1, NCOA6, RXRA, PPARA, PPARG, HELZ2); Uptake and trafficking (LDLR, LPL, CXCL16, FABP5, ABCA2, SAR1B, SEC24A, SEC24C, SEC24D, VCP, MAL2); Fatty Acid Synthesis (FASN, SCD, ELOVL6, FADS2, PCYT2, CHKA); and Indirect Sterol Regulation (LGMN, PDK3, ATF5, CARM1, TP53INP1, FBXO6, S100A11, AVPR1A, KPNB1, TNFRSF12A, GUSB, MTF1, NFIL3, ATF3, PLAUR, TRIB3, PLSCR1, ACTG1, ECH1, ANTXR2, CTNNB1, JAG1, CLU, SEMA3B, TGS1, ALCAM, ERRFI1, TBL1X, ALDOC, GNAI1, CD9, ADH4, ACAT2). After removal of duplicate genes across overlapping gene sets, the resulting panel contained 72 unique cholesterol associated genes. Differential expression results were filtered for this gene set and visualized across pairwise comparisons of

benign nevi versus early stage primary melanoma, benign nevi versus late stage melanoma, and early versus late stage melanoma (q-value < 0.05 was considered significant).

***Survival Curve Analysis***

Survival analysis was performed using the Skin Cutaneous Melanoma TCGA PanCancer Atlas cohort (TCGA-SKCM)[9] accessed through the cBioPortal for Cancer Genomics[10]. For each functional cholesterol homeostasis pathway, genes identified as differentially expressed between benign nevi and early stage primary melanomas in the GSE98394 dataset were used to define the gene set for survival analysis input. Patients in the TCGA-SKCM cohort were grouped into altered or unaltered groups according to mRNA expression, and Kaplan-Meier estimates of overall survival were generated directly in cBioPortal, and the differences between groups were assessed using the log rank test.

### Confinement assays

To reproducibly confine cells during high-resolution live-cell imaging, confinement was performed using the 1-well Dynamic Cell Confiner System (4Dcell) per manufacturer's directions. Briefly, the system consists of a Cobalt autonomous vacuum pump attached to a manufactured PDMS suction cup fitted with a glass coverslip containing PDMS micropillars of 3 µm in height which determine the spacing between the top of the coverslip and the cell culture dish. To perform confinement experiments, cells were transiently transfected and plated for 24-48 hours on 35mm glass bottom dishes (FluoroDish, WPI) coated with 10 µg/ml fibronectin. Prior to imaging, the PDMS suction cup and coverslip were briefly sonicated in 70% ethanol, followed by a 5-minute sonication in PBS (Invitrogen), then placed in cell culture media for 1 hour at room temp to equilibrate the PDMS. To confine the cells, the 35mm glass bottom dish was washed briefly with pre-warmed PBS, then 1 ml of fresh warmed cell culture media was added. The sample was then placed on the microscope and the PDMS suction cup was attached by a low pressure vacuum seal that was sufficient for attachment of the PDMS perimeter base to the glass dish, but insufficient for cellular confinement. Confinement was initiated over 1 minute using a ramp of 10 mbar per second utilizing custom software (4Dcell) to control the vacuum pump.

### Microscopy

***Spinning Disk Confocal Imaging***

Imaging was performed on a Nikon Eclipse Ti2 microscope equipped with a Yokogawa CSU-W1 spinning disk scanhead, a Nikon motorized stage with a Nano-Z100 piezo insert (Mad City, Madison, WI), and either a Plan Apo 60x oil 1.49 NA DIC. Illumination was provided by a Nikon LUNV 6-line laser unit, and images were captured with a Hamamatsu Orca-Flash 4.0 v3 camera. The system was controlled by NIS-Elements software (Nikon). Cells were imaged in 35 mm glass bottom dishes (FluoroDish). Maintenance of sample temperature and humidity was performed by a Tokai Hit stage-top incubator (Tokai Hit).

***Super-resolution Confocal Imaging***

Imaging of confined live cells for super-resolution was performed using a LSM 880 or 890 Zeiss confocal microscope equipped with an Airyscan using a Plan-Apo 63x 1.4 NA oil objective. Airyscan image reconstructions were processed in auto strength mode using ZenBlack software (Version 2.3). Additional analysis was performed in ImageJ (NIH).

***Fluorescence lifetime Imaging Microscopy (FLIM) Imaging***

***Laurdan FLIM***

Fluorescence lifetime imaging microscopy (FLIM) imaging of Laurdan labeled melanocyte and melanoma cells was performed using an upright Leica Stellaris system equipped with a FALCON FLIM module with a Plan-Apo 60x 1.4 NA objective. The Laurdan fluorophore was excited at 780 nm and emission was collected using two spectrally tunable HyD detectors; their range was set as follows HyD-NDD1 (420 nm-465 nm) for blue emission, and HyD-NDD2 (510 nm-560 nm) for green emission. Spectral and FLIM imaging was performed simultaneously at a frame size of 512 x 512 pixels (pixel sizes of 368 nm in X and Y), with a scan speed of 600 Hz. Laser power was held constant between experimental conditions and days at ~2.4 mW at the back aperture of the objective lens. Line repetition was set to 16 to accumulate sufficient photons for analysis and to maximize collection of photons for detection of the nuclear envelope and to prevent spectral overlap.

***Flipper-TR FLIM***

FLIM imaging of Flipper-TR-labeled cells was performed on a Nikon AX R confocal microscope equipped with a Nikon Spatial Array Confocal (NSPARC) detector and a PicoQuant FLIM system, Plan Apo 60x oil 1.4 NA, Nikon motorized stage with a Nano-Z100 piezo insert (Mad City), and a stage top incubator system (Okolab). Fluorescence lifetime measurements were acquired using PicoQuant time-correlated single-photon counting (TCSPC) system integrated within NIS-Elements software. Cells were labeled with Flipper-TR as described above and was excited using a pulsed 485 nm laser at 20 MHz. The fluorescence emission was collected through a 600/50nm bandpass filter using the PicoQuant FLIM hybrid detector. Images were acquired until a minimum photon count of 100 photons/pixel was reached within our region of interest. Acquisition parameters were kept constant across experimental conditions.

### Image analysis

***BODIPY-cholesterol analysis***

BD-Chol fluorescence was quantified in FIJI (ImageJ, Version 1.54) using segmented membrane regions identified from mCherry-Sec61 using a 10 µm 3 pixel line scan ROI. The nuclear membrane (NM) was defined as the membrane signal surrounding the SiR-DNA labeled nucleus and overlapping with mCherry-Sec61, whereas ER regions were selected from Sec61-positive membranes adjacent to but spatially separated from the nuclear periphery. Plasma membrane (PM) regions were selected along the cell periphery. Because the inner and outer nuclear membranes cannot be independently resolved at this resolution, fluorescence measured at the nuclear periphery does not distinguish the INM from ONM and is the combined NM measurement. To compare relative BD-Chol distribution between membrane compartments while accounting for cellular variation in probe uptake and labeling, mean BD-Chol fluorescence within the ER and NM was normalized to mean PM fluorescence measured within the same cell. The PM was selected as an internal reference based on its established high cholesterol content. PM normalized measurements represent the relative distribution of BD-Chol between membrane compartments and were not used to infer absolute cholesterol abundance. For regional comparisons within the ER,NE and nuclear blebs, BD-Chol fluorescence was normalized to the integrated total cell BD-Chol fluorescence intensity measured within the same cell to account for differences in overall probe labeling. For analysis of cholesterol enrichment at nuclear blebs, BD-Chol fluorescence was measured at the base/neck of individual nuclear blebs and at an adjacent non-bleb region of the NM using equivalent ROIs. Bleb and non-bleb measurements were normalized to integrated total-cell BD-Chol fluorescence and compared pairwise within individual cells. To determine the spatial organization of cholesterol within the NM, BD-Chol fluorescence intensity was sampled along the nuclear periphery using a 10 µm 3 pixel line scan ROI and fluorescence variance was calculated from the intensity values obtained along each NM trace, with higher variance representing increased spatial heterogeneity of BD-Chol fluorescence along the NM.

***Laurdan FLIM measurements***

FLIM data was acquired using the Leica FLIM module as described above. Using the phasor plot approach, the fluorescence decay was acquired at every pixel of the image and a phasor transformation is applied and plotted using the (G,S) coordinates to determine location of each pixel in phasor space[11]. First, with custom Leica software, each Leica FLIM data file was converted to both a summed intensity image, used for segmentation of subcellular structures, and a stack of 139 longitudinal time bins of width 97 ps each containing photon counts in each pixel. ImageJ was used to isolate different cellular components using the following approaches: whole cell segmentation was performed using the entire image thresholded by intensity; plasma membrane and nuclear envelope, a 5 pixel free hand line was used to segment either the outside edge of all cells in a view field (plasma membrane) or around visible perinuclear regions (nuclear envelope), or a freehand ROI in the internal part of the cell avoiding regions enriched in lipid droplets (intracellular membranes). Following segmentation, the background was removed and segmented ROIs were applied to the time bin pixel image from each emission channel. Following this, a custom built software, written in the IDL programming language (NV5 Geospatial Software (Broomfield, CO)) was used to sum all pixels from each segmented field of view (FOV) and perform a phasor transformation, identifying the peak of the histogram, defined as the center of mass from each image. Each segmented FOV generated a single phasor center-of-mass measurement and therefore constituted one experimental data point. The time base for phasor calculations was determined from the synthetic instrument response function (IRF) included in each Leica data file. By convention we selected the time base to start at one bin back from the peak of the IRF signal to approximate the 2/3 timepoint of the response following each pulsed laser flash. Datasets containing less than 100 photons per pixel, or exhibiting a skewed distribution were excluded from analysis. The angle position of the center of mass of the 3D histogram of pixel density was then determined using the arc tan of its (g,s) coordinates on the phasor plot, and represents the center of mass of FLIM values, and was used to measure the population shift in phasor space due to membrane composition. The custom IDL code contained modules for density plotting from the open source Coyote Graphics Routines Library (http://www.idlcoyote.com/) and are available upon request.

#### *Flipper-TR Image Analysis*
To quantify Flipper-TR fluorescence lifetime at the NM, cells were labeled and imaged with Flipper-TR as described above. Flipper-TR fluorescence lifetime images were analyzed using the Picoquant fast lifetime calculation, which utilizes the mean photon arrival time to estimate fluorescence lifetime per pixel. The NM was segmented by tracing the nuclear periphery using transient expression of mCherry-Sec61 with a 3 pixel wide line ROI, this ROI was then applied to the average lifetime FLIM channel and measurements were exported to GraphPad Prism for analysis. Mean Flipper-TR lifetime was calculated from all pixels contained within the NM ROI for each cell and used as a relative measure of NM tension induced by local lipid packing and composition. For analysis of the relationship between NM tension and NE rupture susceptibility, mean Flipper-TR lifetime for each experimental condition was compared with the corresponding previously published NE rupture[2]. The relationship between mean NM Flipper-TR lifetime and NE rupture frequency was assessed by simple linear regression in GraphPad Prism, with goodness of fit reported as $R^2$.

#### *Point Curvature Analysis*
Nuclear membrane curvature was quantified from mCherry-Sec61 fluorescence using the Kappa curvature-analysis plugin in ImageJ[12]. The Sec61-defined nuclear membrane was traced with a 3 pixel line scan for each nucleus and absolute point curvature was calculated along the resulting membrane contour using an open trace and reported in $\mu m^{-1}$. Nuclear blebs were traced and analyzed separately from non-bleb regions of the nuclear membrane. To determine how confinement and LBR depletion altered the distribution of nuclear membrane curvature, point-curvature measurements were analyzed independently for unconfined nuclear membrane, confined nuclear membrane, confined nuclear blebs, and confined LBR-depleted cells (n=10 cells per condition). For each cell, point-curvature measurements were assigned to predefined curvature intervals of 0–0.05, 0.05–0.10, 0.10–0.15, 0.15–0.20, 0.20–0.25, 0.25–0.30, 0.30–

0.35, 0.35–0.40, 0.40–0.45, 0.45–0.50, 0.50–1, 1–2, 2–5, 5–10, 10–20, 20–50, and ≥50 $\mu m^{-1}$. The number of measurements within each curvature interval was expressed as a percentage of the total number of point-curvature measurements obtained from the same cell. Curvature distributions were therefore calculated independently for each nucleus before comparison between experimental conditions, with individual cells treated as biological replicates.

*Analysis of fluorescence intensity and local membrane curvature*

To determine the relationship between local cholesterol enrichment and nuclear membrane curvature, matched point-curvature and fluorescence-intensity measurements were obtained along the same membrane contour. Analyses were performed independently for each nucleus to account for differences in absolute fluorescence intensity between cells. Within each cell, membrane positions were ranked according to fluorescence intensity and divided into 10 percentile intervals spanning 0-100% fluorescence intensity. Mean absolute point curvature was then calculated for all membrane positions contained within each fluorescence percentile interval. Percentile intervals were represented by their midpoint values (5, 15, 25 ... 95%) for graphical analysis. Individual cells were maintained as independent biological replicates and raw point measurements were not pooled between cells. The same analysis was performed independently using BD-Chol and mCherry-Sec61 fluorescence. For BD-Chol, analyses were additionally performed separately for nuclear bleb and non-bleb nuclear membrane regions to determine whether the relationship between cholesterol enrichment and curvature differed between membrane regions. mCherry-Sec61 was analyzed using the same approach to determine the relationship between membrane-associated fluorescence and local membrane curvature independently of BD-Chol enrichment. To determine whether regions of highest cholesterol enrichment preferentially localized to nuclear blebs, BD-Chol fluorescence measurements within each nucleus were ranked by intensity and values at or above the 95th percentile were identified. The relative distribution of these measurements within nuclear bleb and non-bleb NM regions was quantified within individual nuclei and compared pairwise.

***Statistical analysis***

Statistical analyses were performed using GraphPad Prism (version 9). The experimental n represents individual cells/FOV/patients, and N represents independent biological experiments are indicated in the corresponding figure legends. Data are presented as mean ± SD or mean ± SEM as indicated. Pairwise measurements obtained from different regions within the same cell were analyzed using paired statistical tests, whereas comparisons between independent experimental groups were analyzed using unpaired tests. Comparisons involving more than two groups were performed using one-way ANOVA followed by multiple-comparison tests. Two-group comparisons were performed using Student's t test with Welch's correction or Mann-Whitney as indicated in the legends. Relationships between continuous variables were assessed using simple linear regression. Statistical significance was defined as*p ≤ 0.05, ** p ≤ 0.01, ***p ≤ 0.001, and ****p ≤ 0.0001, n.s is not significant.

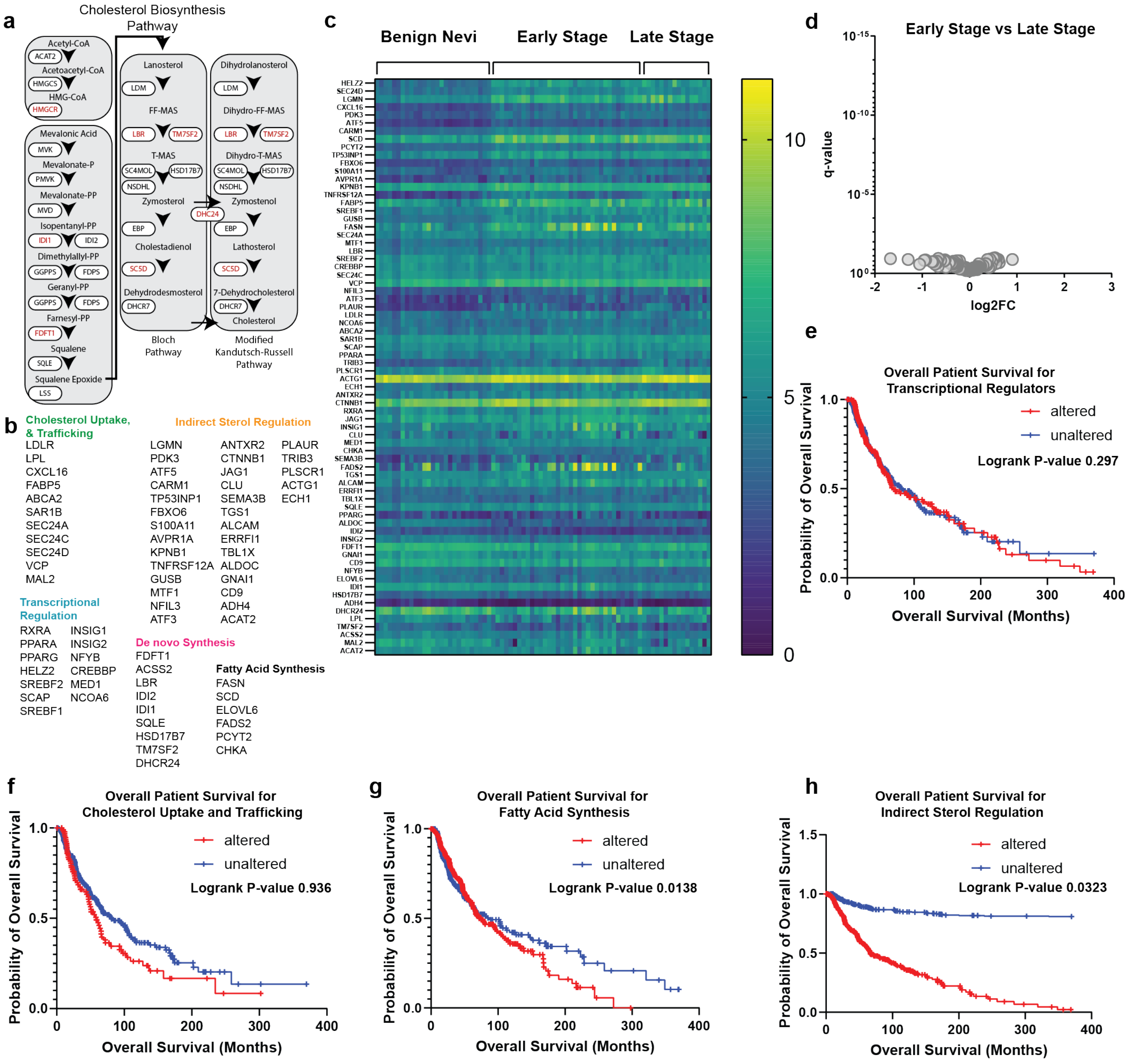


**Figure S1. (A)** Schematic of the de novo cholesterol biosynthetic pathway from acetyl-CoA to cholesterol. Major sterol intermediates and enzymes are indicated, including the alternative Bloch and modified Kandutsch-Russell pathways. Enzymes in red indicate enriched genes from our analysis. **(B)** List of genes in each cholesterol functional pathway**. (C)** Heat map of transcript abundance for the complete curated set of 72 cholesterol-associated genes across individual benign nevi, early stage primary melanomas, and late stage primary melanomas from (GEO:GSE98394). **(D)** Volcano plot of differential expression of the 72 cholesterol-associated genes comparing early with late stage primary melanomas. Differential expression is plotted as log2 fold change (log2FC) versus q-value. All genes were non-significant genes with a q>0.05 (grey). **(E–H)** Kaplan-Meier analysis of overall survival in patients from The Cancer Genome Atlas (TCGA) PanCancer melanoma cohort (SKCM) stratified by altered or unaltered expression of genes associated with (D) Transcriptional Regulators (13 Genes; N=425 patients; Altered group=235, Unaltered group=190) Log rank P-value=0.297, (E) Cholesterol Uptake and Trafficking (11 Genes N=425 patients; Altered group=194, Unaltered group=231) Log rank P-value=0.936, (F) Fatty Acid Synthesis (6 Genes; N=426 patients; Altered group=104, Unaltered group=322) Log rank P-value=0.0138 and (G) Indirect Sterol Regulation (33 Genes; N=426 patients; Altered group=326, Unaltered group=100) Log rank P-value=0.0323. Gene sets used for survival analysis were limited to genes within each functional cholesterol homeostasis category that were differentially expressed between benign nevi and early primary melanomas. In (C), q ≤ 0.05 was used to determine significance; in (E-H), significance was determined by log-rank test.

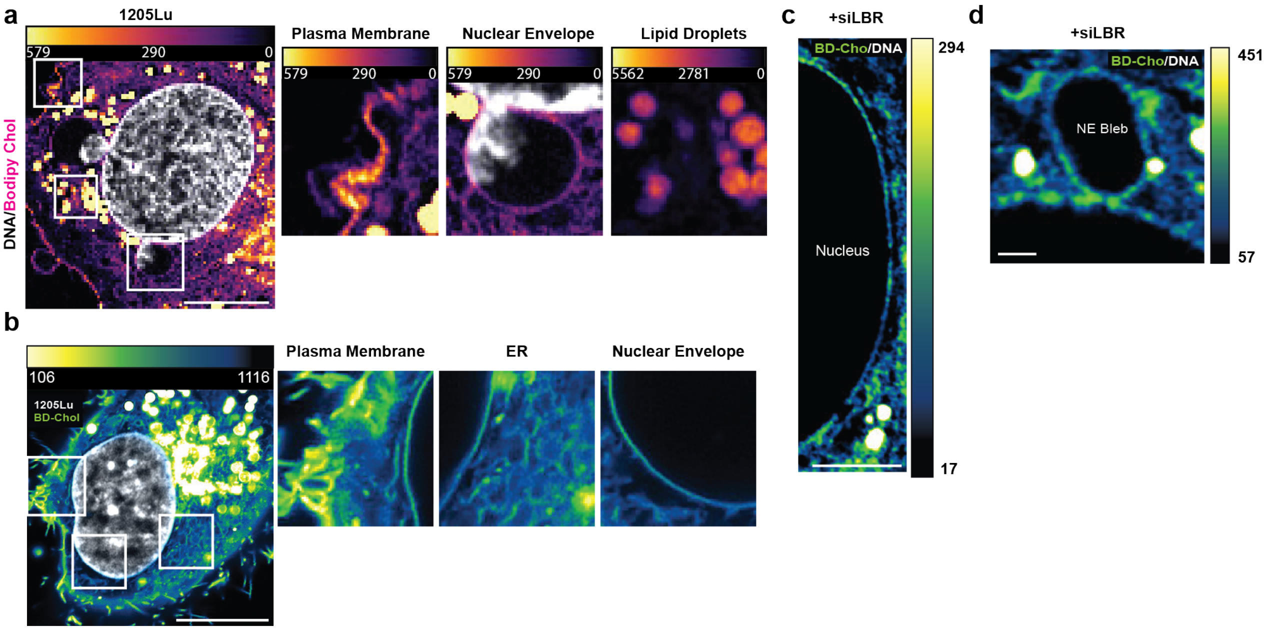


**Figure S2. (A)** Intensity color-coded super-resolution image taken during confinement to 3 µm of 1205Lu melanoma cell labeled with BODIPY-Cholesterol (BD-Chol) and SiR-DNA (Grey) to visualize cholesterol and chromatin. Boxes indicate subcellular regions enlarged at right representing examples of BD-Chol localization at the plasma membrane, nuclear envelope, and lipid droplets. **(B)** Intensity color-coded super-resolution image taken of an unconfined 1205Lu cell labeled with BODIPY-Cholesterol (BD-Chol) and SiR-DNA (Grey) to visualize cholesterol and chromatin. Boxes indicate subcellular regions enlarged at right representing examples of BD-Chol localization at the plasma membrane, endoplasmic reticulum (ER), and nuclear envelope. **(C)** Higher contrast intensity color-coded super-resolution image of the nuclear envelope in an LBR-depleted (+siLBR) 1205Lu cell from (Fig 2F). **(D)** Higher contrast intensity color-coded super-resolution image of a nuclear bleb in an LBR-depleted (+siLBR) 1205Lu cell from (Fig 2I). In (A, B) bar = 10 µm, in (C) bar= 5 µm, (D) bar= 1 µm.

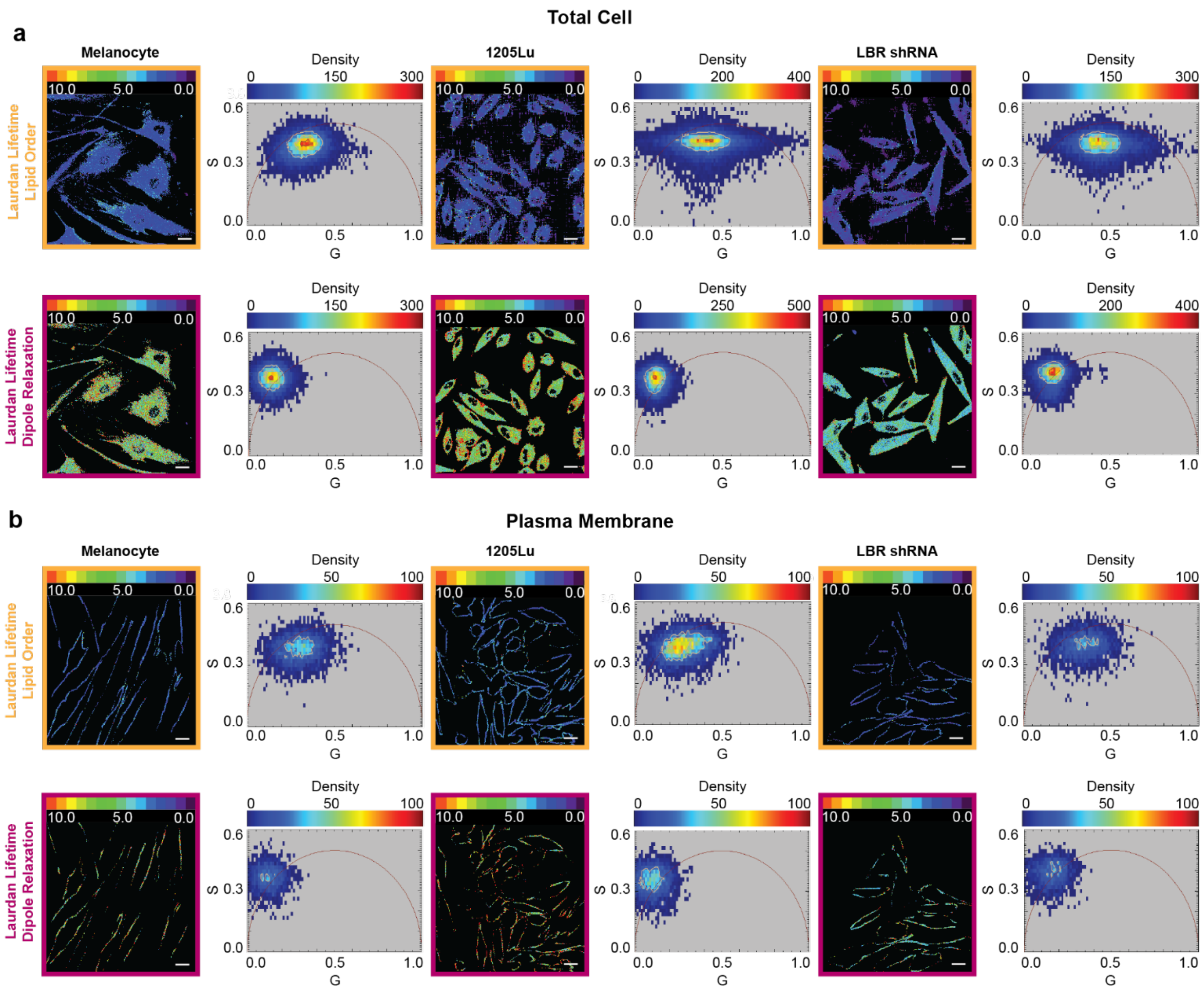


**Figure S3. (A)** Representative two-photon FLIM images (right panels) of melanocytes, 1205Lu and LBR shRNA cells in a blue emission (lipid order; Orange Boxes) or green emission (dipole relaxation; Magenta Boxes) and their corresponding phasor plots (left panels). **(B)** Representative segmentation images of plasma membrane with corresponding phasor plot. All experiments were performed using lipid depleted media. Color scale bars over images represent FLIM lifetime values, over phasor plots represent pixel density. Scale bar = 10 μm.

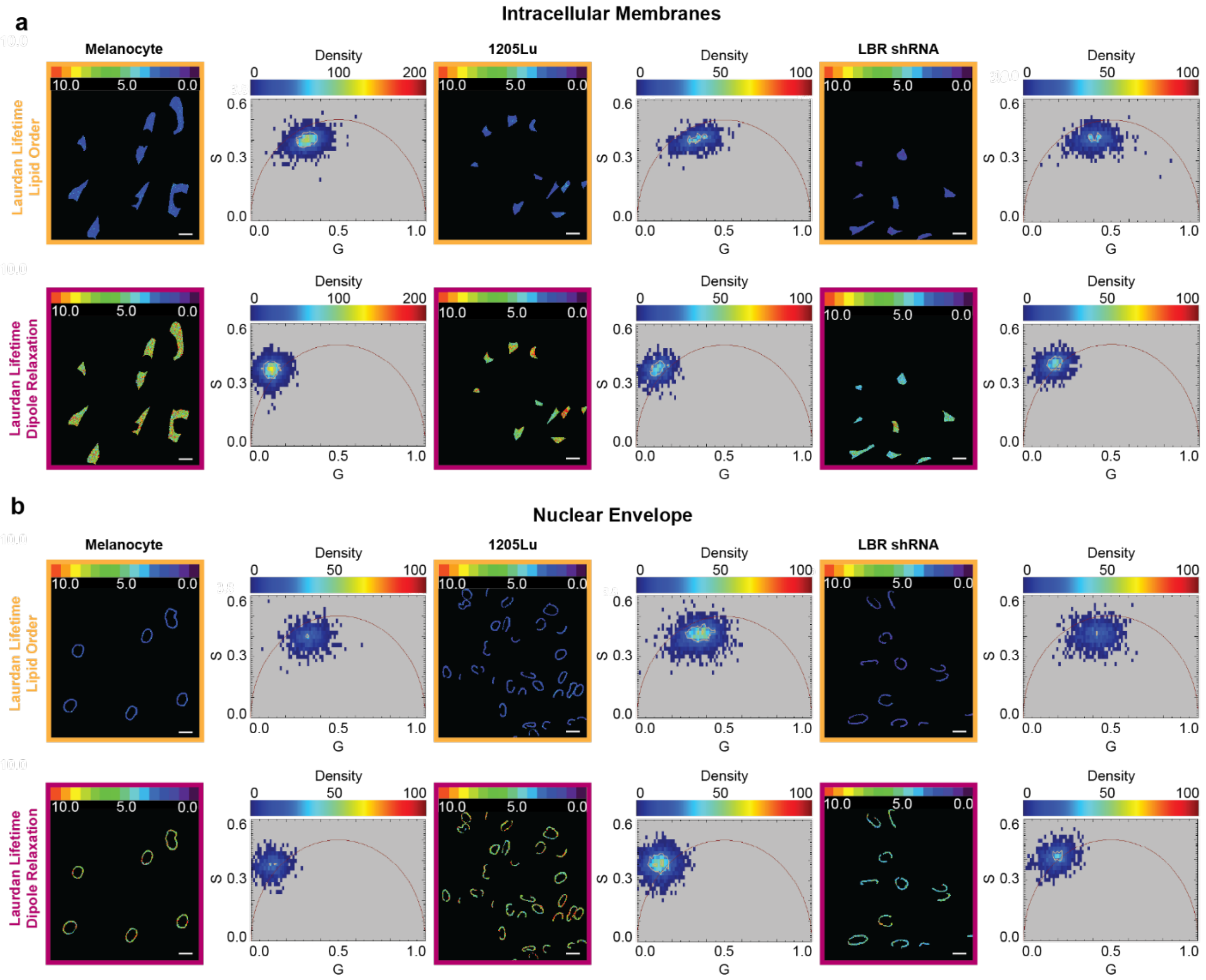


**Figure S4. (A)** Representative FLIM images of melanocytes, 1205Lu and LBR shRNA cells in a blue emission (lipid order; Orange Boxes) or green emission (dipole relaxation; Magenta Boxes) and their corresponding phasor plots depicting pixel density segmented internal membranes (A), or the nuclear envelope **(B)** with corresponding phasor plot. All experiments were performed using lipid depleted media. Color scale bars over images represent FLIM lifetime values, over phasor plots represent pixel density. Scale bar = 10 µm.

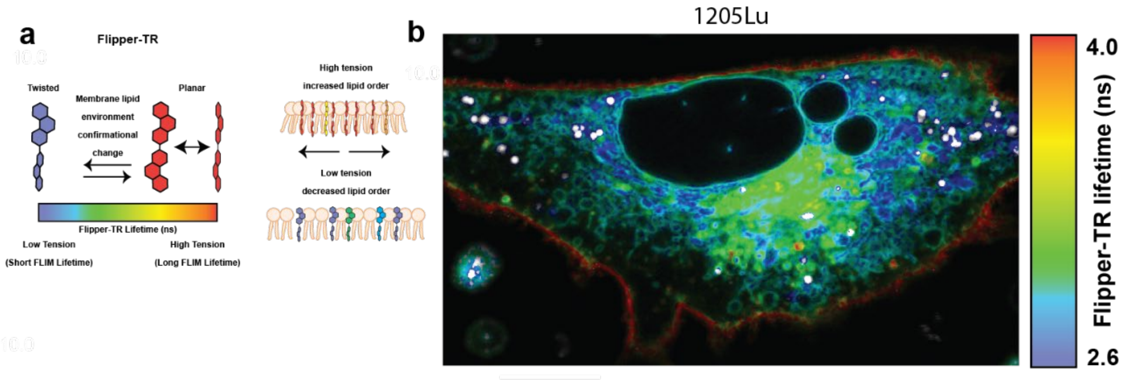


**Figure S5. (A)** Schematic of Flipper-TR, a mechanosensitive fluorescent probe whose planarization within the membrane lipid bilayer due to lipid composition and increased lipid packing induced by membrane tension increases fluorescence lifetime. **(B)** Intensity color-coded FLIM image of 1205Lu melanoma cell labeled with Flipper-TR. Scale bar =  10 μm.
